# EXPERIMENTAL CHARACTERIZATION OF BALLOON ANGIOPLASTY IN HUMAN FEMOROPOPLITEAL ARTERIES WITH DIFFERENT CALCIUM BURDENS

**DOI:** 10.1101/2025.11.27.691029

**Authors:** Eric Anttila, Majid Jadidi, Kaspars Maleckis, Paul Aylward, Anastasia Desyatova, Jason MacTaggart, Alexey Kamenskiy

## Abstract

**Introduction:** Balloon angioplasty is one of the most common treatments for Peripheral Artery Disease (PAD), but its clinical outcomes continue to disappoint, particularly when managing calcified lesions. We characterized luminal gains and damage after balloon angioplasty using high-resolution imaging, histology, and mechanical testing.

**Methods:** Fresh diseased human femoropopliteal arteries (FPA) from n=15 subjects (average age 69 ± 9, range 53-90 years) with different calcium burdens were imaged before, during, and after angioplasty using micro-computed tomography, and luminal gains, calcium fractures, and resulting dissections were quantified. Histology was used to assess structural damage, and biaxial mechanical testing determined damage initiation stretches and stresses.

**Results:** In severely calcified FPAs, calcification often manifested as rings or large plate-like deposits. When calcium spanned the entire circumference, angioplasty produced longitudinal cracks but ∼8% luminal area gain. In less calcified arteries, damage manifested primarily as tears along the internal elastic lamina and within the tunica media. Dissections were present in 53% of all FPAs after angioplasty, with a higher prevalence in more calcified vessels and vessels with stenosis (75% each). Damage initiated at lower biaxial stretches (1.11 ± 0.02 vs 1.14-1.15) and lower longitudinal stresses (44 ± 22 kPa vs 57 ± 31 kPa) in severely calcified specimens compared with lightly calcified arteries, but larger calcium burdens generally required more circumferential stress to initiate damage (66 ± 27 kPa). Diabetes mellitus was associated with a higher calcium burden.

**Conclusions:** Severe calcification limits luminal gains after angioplasty. Less calcified arteries accumulate damage to the healthier wall while calcium remains mostly intact. These results may inform clinical strategies and the development of better devices to treat PAD.

## 1. INTRODUCTION

Peripheral artery disease (PAD) commonly manifests as an atherosclerotic narrowing of the femoropopliteal artery (FPA) lumen that obstructs blood flow^1^. Patients suffering from PAD are at higher risk for major cardiovascular events, including heart failure, myocardial infarction, ischemic stroke, and sudden cardiac death^2–5^, and the economic burden of PAD on healthcare exceeds $21 billion in the United States alone^6^. Failed interventions requiring repeat procedures are a major contributor to this cost, and although the exact reasons for poor clinical outcomes of PAD repair are not completely understood, the interaction between the repair device and the diseased artery likely plays a crucial role^7–9^.

Endovascular intervention using balloon angioplasty, often followed by stenting, is one of the most common treatments for advanced PAD lesions^10^. During balloon angioplasty, a balloon-tipped catheter is percutaneously inserted into the common femoral artery (or less commonly into the brachial or pedal artery) and navigated over a thin guidewire to the area of obstruction using fluoroscopy. The balloon is then inflated, crushing the obstruction and stretching the arterial wall, thereby widening the lumen to increase blood flow to the distal tissues. While angioplasty alone may sometimes be sufficient to resolve the occlusion, flow-limiting arterial dissections due to delamination of the plaque are not uncommon^11^. They are particularly prevalent when treating complex lesions^12^, which often requires the use of a stent.

Previous studies have examined balloon angioplasty results in a clinical setting, often comparing it to other types of surgical repair^13–15^, evaluating drug-coated and uncoated balloons^16–19^, analyzing the effects of inflation time^20^ and balloon length^21,22^ on clinical outcomes, or developing computational models of balloon angioplasty to study device-artery interactions^23^. Nevertheless, the question of what exactly happens to the lesion and to the adjacent healthier wall during angioplasty remains poorly understood. In this study, we have attempted to answer this question and test the hypothesis that luminal gain from angioplasty is directly dependent on calcium burden. To achieve this, we have quantified luminal gains, damage, and dissections resulting from balloon angioplasty under controlled conditions using high-resolution computerized tomography (CT) imaging, histology, and biaxial mechanical evaluation of *ex vivo* diseased human FPAs. These results provide a better understanding of the reasons behind poor clinical outcomes of endovascular PAD repair, and could inform treatment strategies when dealing with complex calcified lesions. They can also help improve computational models of balloon angioplasty aimed at studying device-artery interactions by making them more realistic.

## 2. MATERIALS AND METHODS

### 2.1 Materials

#### 2.1.1 Arterial specimens

Fresh FPAs from 15 subjects 53-90 years old (average age 69 ± 9 years) were obtained from an organ procurement organization, Live On Nebraska within 24 hours of death after receiving consent from next of kin. Subjects were predominantly male (87%), 73% had hypertension (HTN), 47% diabetes mellitus (DM), 40% dyslipidemia (DYS), 53% coronary artery disease (CAD), and 93% were current or former tobacco users (average pack-year history 46 ± 41).

#### 2.1.2 Micro-computed tomography (micro-CT) sample preparation

FPAs were freed from the surrounding connective tissue, muscle, and fat and inspected for regions of gross pathology. The most diseased segments were isolated into approximately 4 cm long sections, and side branches were ligated. The artery was then sutured onto a custom 3D printed setup (**Figure 1**) designed in SolidWorks (Dassault Systèmes, Vélizy-Villacoublay, France) that allowed for the specimen to be pressurized to 80 mmHg. This setup also allowed the insertion and removal of a percutaneous transluminal angioplasty balloon and restricted the movement of the FPA during scanning. Pressurization of arteries was done using aerosol-based shaving foam because it has a significantly lower attenuation factor than the artery itself and allowed the vessel wall to be visible on micro-CT.

### 2.2 Micro-CT scanning and the analysis of pre, during, and post-angioplasty results

FPA segments were scanned perpendicular to the FPA longitudinal axis using a Bruker Skyscan 1172 micro-CT (Bruker Corporation, Billerica, MA). All specimens were scanned before, during, and after angioplasty. Before scanning the artery with the inflated balloon, the shaving foam was flushed out using 0.9% phosphate-buffered saline (PBS) introduced via a 10 cc syringe. Boston Scientific Charger OTW (Boston Scientific Corporation, Marlborough, MA, USA) angioplasty balloons were chosen based on the arterial lumen diameter measured using the pre-angioplasty scan, and the oversizing was approximately 1 mm. The balloon was introduced into the artery via the custom 3D printed setup and inflated with mineral oil to the manufacturer-recommended nominal balloon pressure of 12 atm. Mineral oil was used because it has a lower attenuation coefficient than the artery wall and is non-toxic. The balloon was sutured twice with an ultra-strength fishing line to ensure no pressure leaks, and the catheter immediately proximal to the balloon was cut to allow the setup to fit within the micro-CT device. After scanning the artery during angioplasty, the balloon was deflated and removed, and the FPA was pressurized again to 80 mmHg with shaving foam for the final post-angioplasty scan.

Source voltage and current were set to 44 kV and 226 µA for all three scans, respectively. Other scan parameters included a rotational step of 0.5°, a 0.5 mm Al filter, and 26.94 µm isotropic resolution. The duration of each of the three scans was 28 minutes. Raw rotational TIFF images were converted to cross-sectional bitmap images and reconstructed using NRecon (Micro Photonics, Inc., Allentown, PA). Cross-sectional bitmap images were then exported from NRecon and stitched together in Materialise Mimics (Materialise NV, Leuven, Belgium). Soft tissue and calcification were thresholded based on Hounsfield units, with values >100 generally representing calcification and values ≤100 representing soft tissue. 3D models were created for each sample with soft tissue and calcification to visualize the effects of angioplasty. Soft tissue and calcification surface areas were quantified using Mimics.

Following 3D model reconstruction, calcification burden percentage by volume was quantified for each specimen using Mimics. The average burden across all samples was 13±10% with a maximum of 28% and a minimum <1%. Samples were then classified into one of two groups: severe or mild calcification. Severely calcified arteries (*n* = 8) contained large ringed and plate-like calcium deposits that spanned the majority (>180°) of arterial circumference and constituted 16%-28% of the volume. Mildly calcified vessels (*n* = 7) had <1%-7% of calcium by volume, and contained ringed and plate-like structures that spanned less than half (<180º) of the arterial circumference and were smaller and more localized in nature. In order to quantify the results of angioplasty, we have calculated luminal gain as the change in luminal stenosis for specimens that demonstrated a stenosis that would likely require angioplasty in a clinical setting. Specifically, open lumen area was assessed before and after angioplasty every 269.4 µm along the length of the specimen and averaged in Mimics, resulting in:

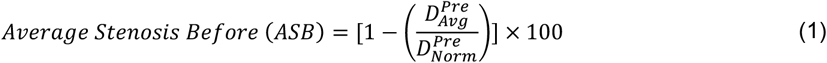

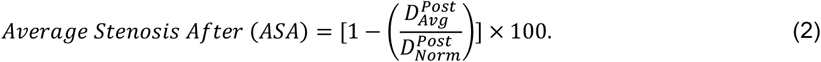

Here *D*_*Avg*_ is the luminal diameter calculated from the measured area and averaged over the length of the artery, and *D*_*Norm*_ is the “healthy” luminal diameter measured at the location of the largest luminal area of the scanned region. Percent luminal gain was then calculated as:

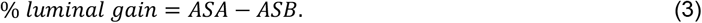

Maximum stenosis before and after angioplasty was quantified using equations:

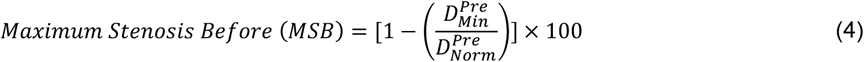

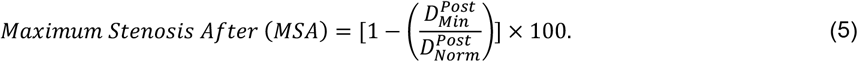

Here *D*_*Min*_ corresponds to the diameter of the smallest area along the length of the sample, while *D*_*Norm*_ is the “healthy” luminal diameter measured at the location of the largest luminal area of the scanned region.

Luminal area along the length of all specimens was quantified before and after angioplasty in order to visualize what was occurring to each sample as a result of angioplasty. 3D reconstructions of the soft tissue and calcium were exported from Mimics as stereolithography files, and a kinematic framework was used to “unwrap” the tubular artery into a flat sheet^24^, thereby allowing easier visualization and analysis of changes after angioplasty. This involved a custom-written image analysis script in MATLAB (MathWorks, Natick, MA) that automatically segmented calcification. Intramural structural changes associated with balloon damage were studied using methacarn-fixed and Verhoeff-Van Gieson (VVG)-stained transverse arterial sections. Four circumferential rings from the angioplasty segment were compared to the section of the adjacent artery that was used as a control (Figure 1), and the damage to the internal and external elastic lamina and the tunica media was assessed.

**Figure 1.**
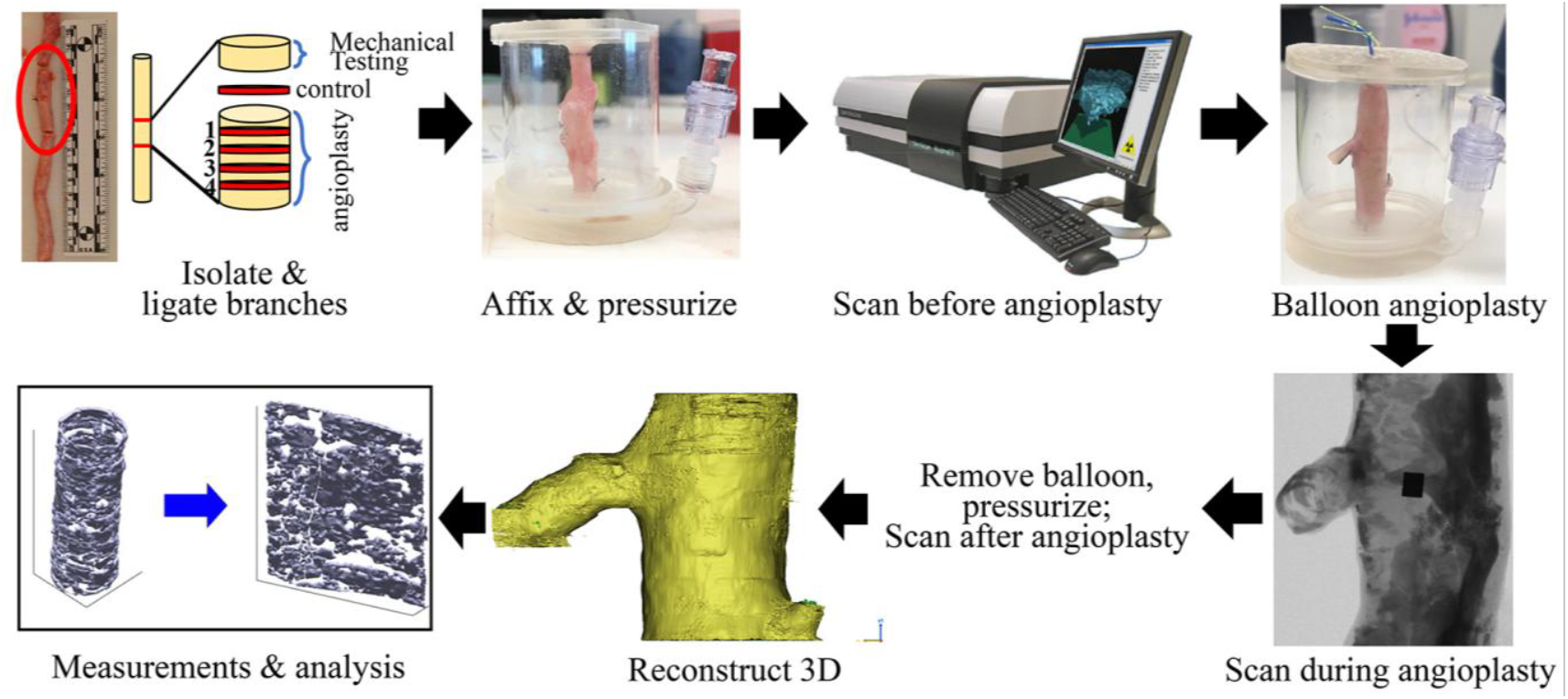
Study flowchart describing specimen preparation, scanning of the artery before, during, and after angioplasty, and the analysis of data.

### 2.3 Mechanical testing

Planar biaxial extension in elastic and inelastic domains was performed on the adjacent arterial sections (Figure 1). These tests were used to assess tissue stiffness and damage initiation stresses and stretches. Experiments were done using 13×13mm specimens radially-opened and submerged in 0.9% PBS at 37^*°C*^. Tests were performed using a CellScale Biotester (CellScale Biomaterials Testing, Waterloo, Ontario, Canada) equipped with 2.5 N loadcells. All samples were first evaluated in the elastic domain starting with 20 equibiaxial cycles of preconditioning to produce a repeatable response and following with 19 multi-ratio loading protocols as described previously^24–26^. Several equibiaxial cycles were mixed in throughout the elastic characterization to ensure the absence of damage. After the elastic characterization, an equibiaxial damage protocol^27^ consisting of 5 cycles of repeated loading and unloading at each stretch limit was executed, and the stretch limits were increased with a step of 0.025-0.05 until the force reached the load cell limit. These tests allowed to determine damage initiation stretches and stresses defined as those at which the preconditioned FPA stopped demonstrating the repeatable elastic response and started exhibiting inelastic stress softening effects^27^. Since large supraphysiologic stretches damaged the tissue, it was not possible to run multi-ratio tests, and all data were obtained from an equibiaxial protocol conducted at 0.01 s^-1^ strain rate^27^.

## 3. RESULTS

Figure 2 demonstrates all analyzed arteries before, during, and after angioplasty organized from the largest to smallest calcium burden and in two views: longitudinal and transverse. Severely calcified specimens (#1-8, red) generally demonstrated large ring- and plate-like calcium deposits that spanned the whole circumference and length of the artery, while sample #4 and #7 had many small calcium nodules dispersed in the wall and in the plaque. All specimens from this group had branch vessels within the scan region with calcification buildup at or near the branches, and five of the eight (63%) specimens had DM. After angioplasty, six of the eight (75%) arteries developed dissections. Mildly calcified specimens (#9-15, green) had primarily small, isolated calcium structures that typically spanned less than half of the arterial circumference. Most of these specimens had luminal stenosis, but the plaque was not heavily calcified. Five of these seven specimens had a branch vessel in the scanned region, and two (29%) developed dissections after angioplasty. Dissections were present in six of eight (75%) stenosed specimens, while two of seven (29%) non-stenosed specimens developed a dissection after angioplasty. The overall rate of dissections after angioplasty across all specimens was 53%.

**Figure 2.**
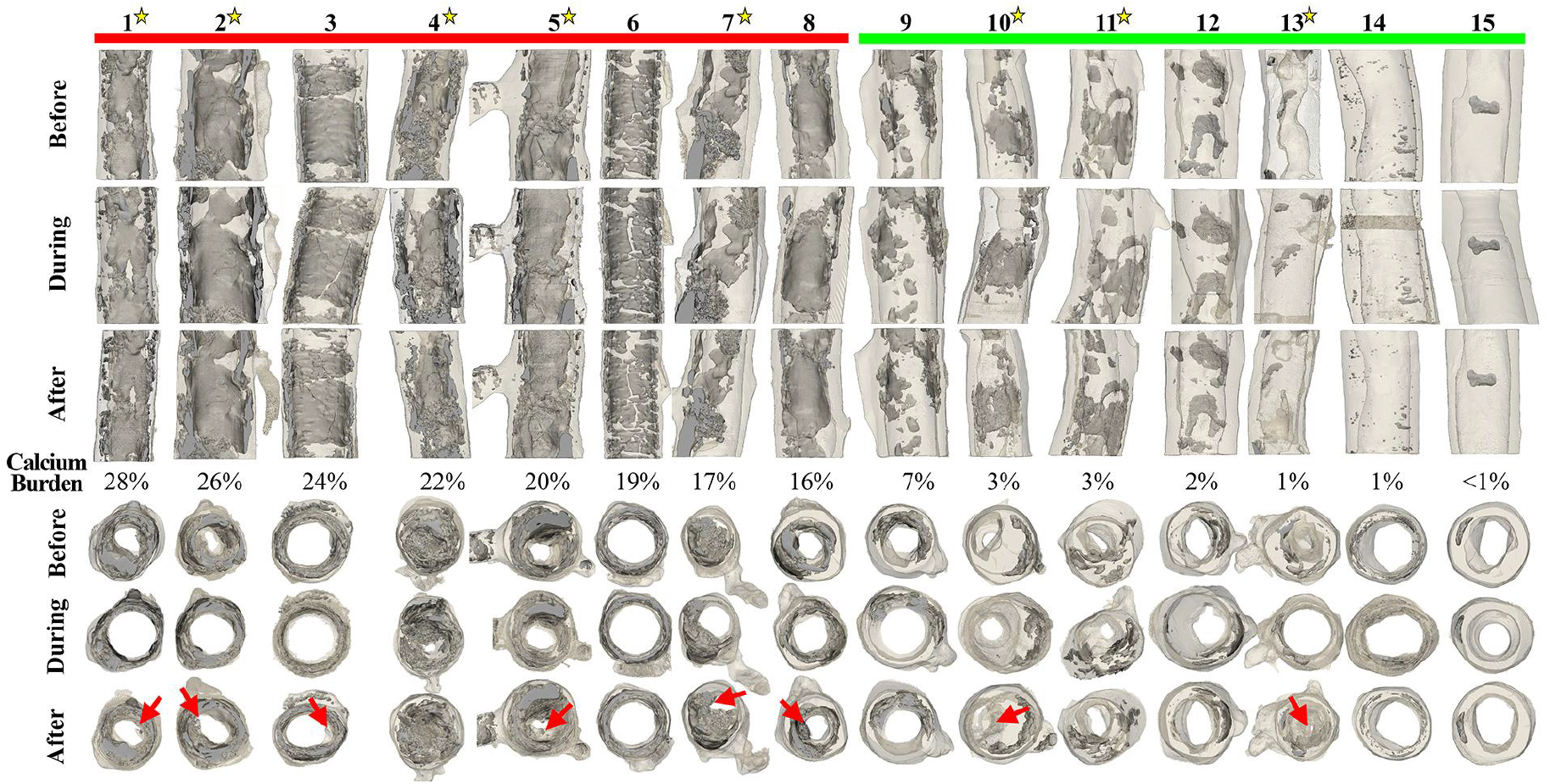
3D reconstructions of all 15 FPA specimens before, during, and after angioplasty. Specimens are arranged left to right from the largest to smallest calcium burden (%). Color bars on top correspond to the two calcification groups: severe (red) and mild (green). Both longitudinal and transverse views are presented, and red arrows on transverse views point to the dissections as a result of angioplasty. Yellow stars next to the sample number correspond to samples that were considered to be stenosed. Luminal gains (%) after angioplasty are summarized in the bottom row.

Luminal area along the axial length before and after angioplasty for all samples is shown in Figure 3. Average and maximum stenosis values are shown for specimens with a substantial stenosis that would likely require angioplasty in a clinical setting. The majority of samples demonstrate regions along the length that does not improve luminal area as a result of angioplasty. Only samples 8, 9, and 14 demonstrated increased luminal area through the entire length of the scanned region. Samples 1, 3, 5, and 6 from the severely calcified group demonstrated multiple regions where luminal area was greater before angioplasty than after it.

**Figure 3.**
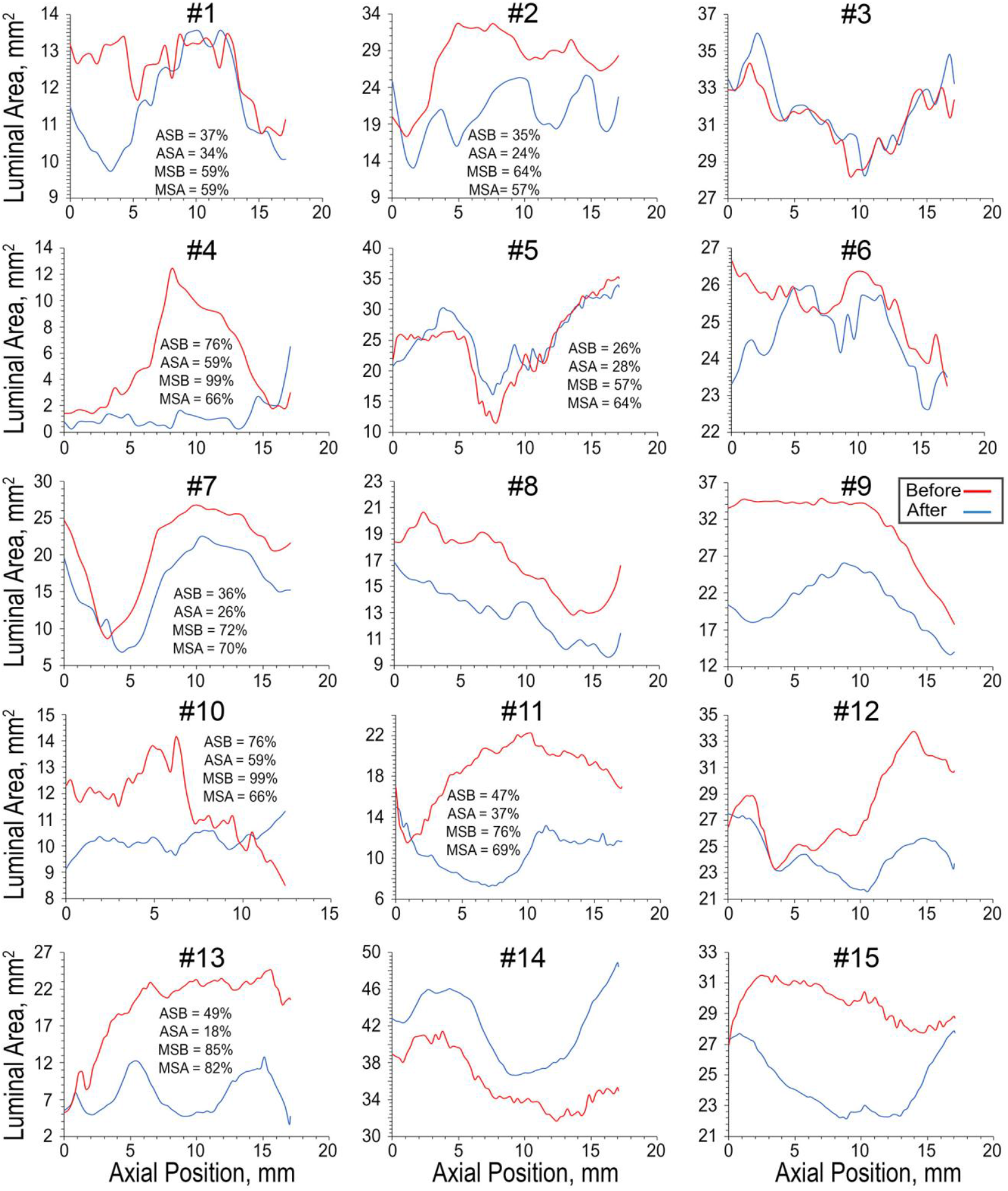
Luminal area versus axial position for all samples before (blue) and after (red) angioplasty. Axial position 0 corresponds to the bottom of the sample. ASB = average stenosis before, ASA = average stenosis after, MSB = maximum stenosis before, MSA = maximum stenosis after.

Figure 4A demonstrates luminal gain as a function of calcium burden for stenosed samples only (Samples #1, 2, 4, 5, 7, 10, 11, 13). Severely calcified specimens had the smallest luminal gains after angioplasty of just 8.0 ± 7.2%. Sample #4 had the highest gain in this group (17%), but the residual stenosis was still 59% after angioplasty. One of the five severely calcified arteries with stenosis showed a negative luminal gain, meaning that the lumen became smaller after angioplasty because of the protruding dissection. Mildly calcified arteries had an average of 14.5 ± 14.1% luminal gains. Sample #13 had the largest gain of all specimens (30%), but it also developed a large dissection (Figure 2). All three samples with stenosis from this group demonstrated a luminal gain as a result of angioplasty. Figure 4B shows the box plot comparison between severe and mildly calcified samples with stenosis.

**Figure 4.**
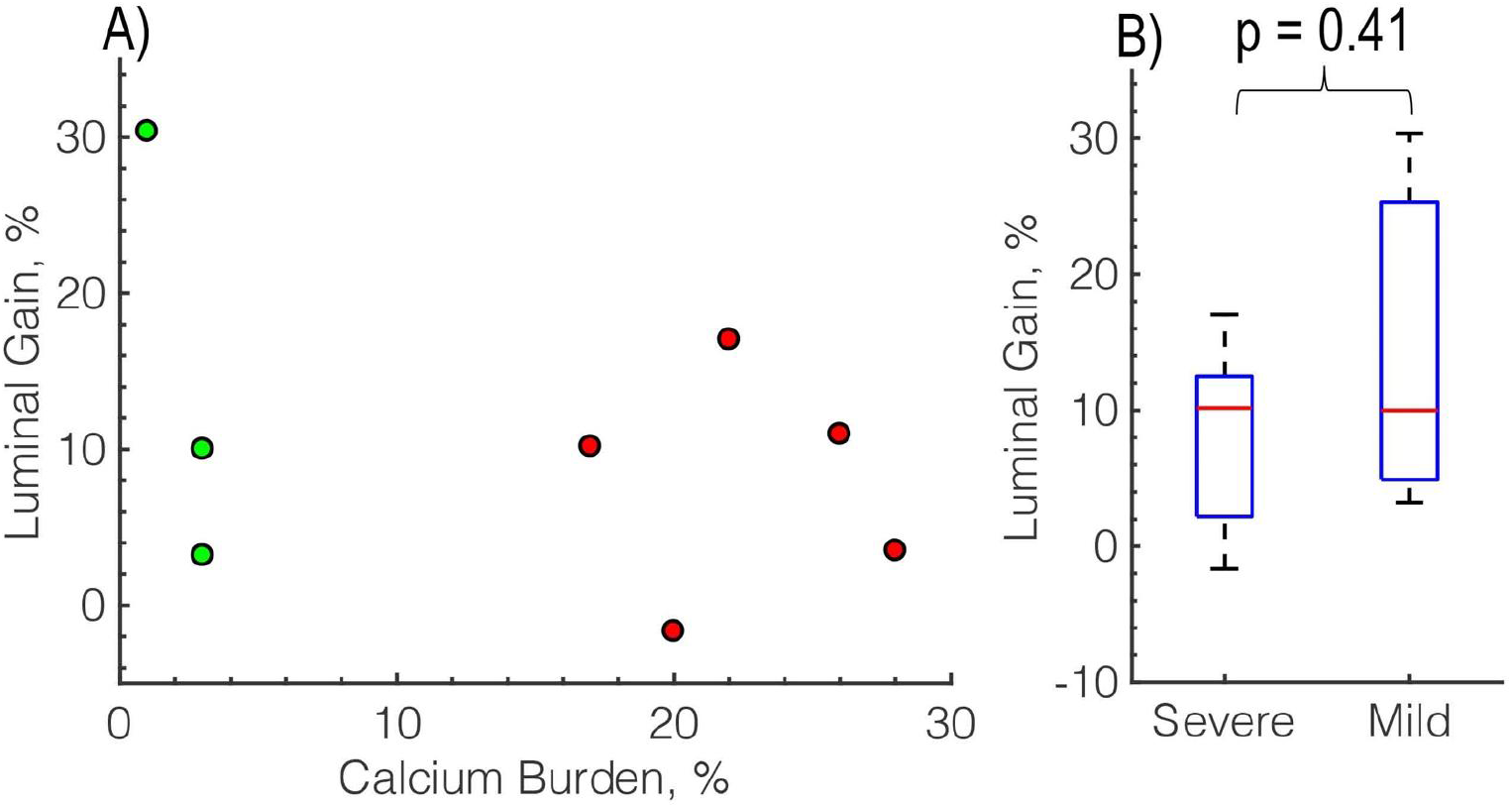
A) Luminal gain after angioplasty (%) plotted as a function of calcium burden (%). The color of the dots corresponds to calcium severity: red = severe, green = mild calcification and B) box plot group comparison of luminal gain after angioplasty (%) in severely (#1, 2, 4, 5, 7) versus mildly (#10, 11, 13) calcified samples with stenosis present.

A closer view of the calcification before, during, and after angioplasty for three representative severely calcified specimens is presented in Figure 5. For better visualization, the circular cross-section is mapped onto the 2D plane, and the black areas of calcification are outlined with red (before), green (during), and blue (after) colors. Red rectangles mark the areas of crack nucleation and growth that occurred primarily in the longitudinal direction. After angioplasty, many of these longitudinal cracks closed back, and the calcium plates returned to their original pre-angioplasty configurations, akin to puzzle pieces. In less severely calcified specimens, little to no evidence of longitudinal cracking was observed, and calcification tended to remain relatively unchanged after angioplasty. Arteries that did not have calcification spanning the entire circumference also had little to no evidence of longitudinal calcium cracking but have instead demonstrated an elongation of the softer tissue next to the calcium deposits (i.e., sample #7 in Figure 2).

**Figure 5.**
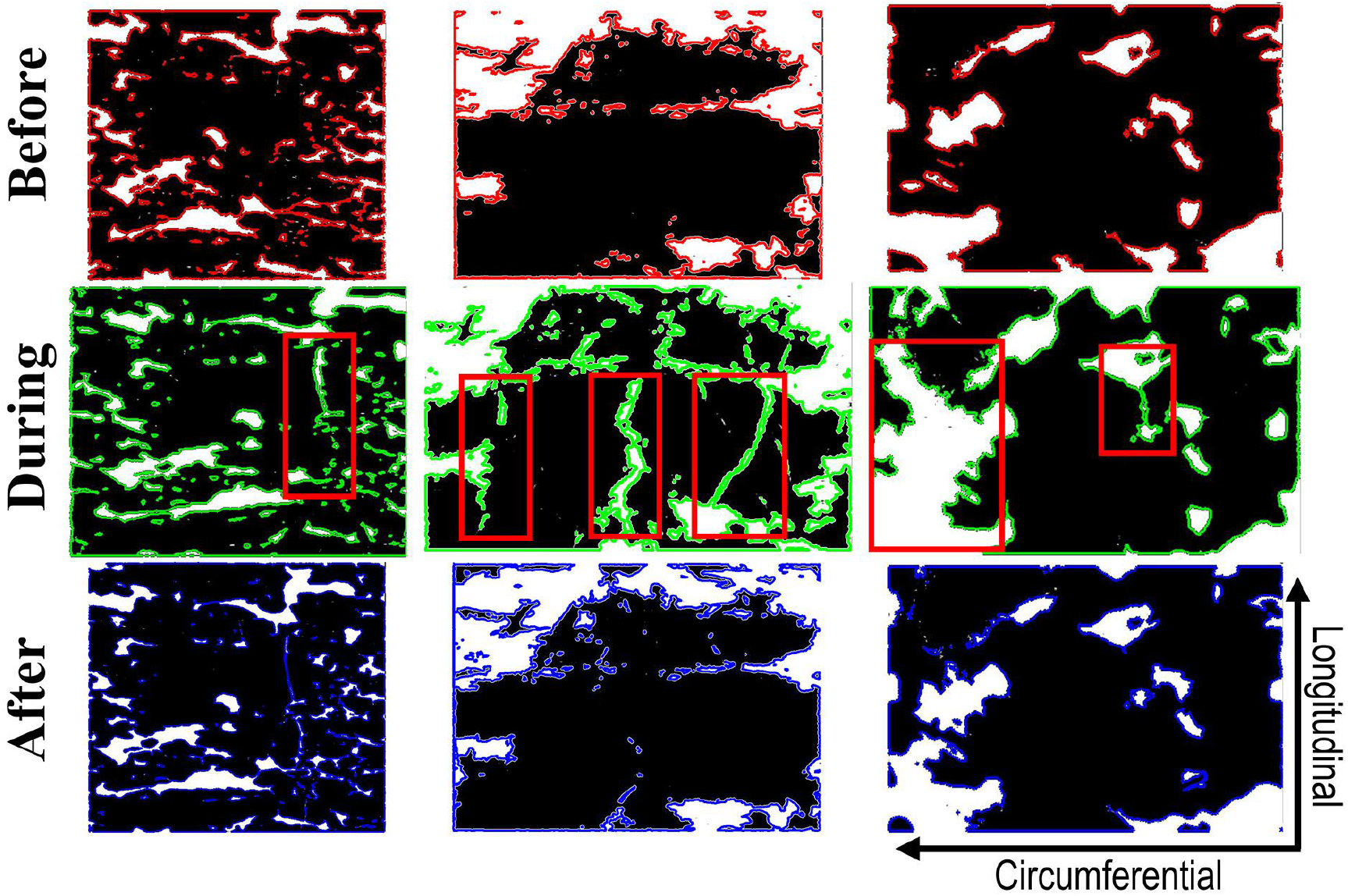
Unwrapped calcification before, during, and after angioplasty for sample #6 (left), #3 (middle), and #2 (right). All three samples were severely calcified. Red rectangles mark the areas of crack nucleation and growth. Calcification is black.

3D views of the calcification in sample #4 before and after angioplasty are depicted in Figure 6. Multiple large longitudinal cracks can be seen as a result of angioplasty, highlighted in the “Front” image. A small pre-crack is present in the “Front” view before angioplasty, although on an adjacent deposit of calcification that appears to not be connected to the large, plate-like structure that experiences substantial cracking as a result of angioplasty.

**Figure 6.**
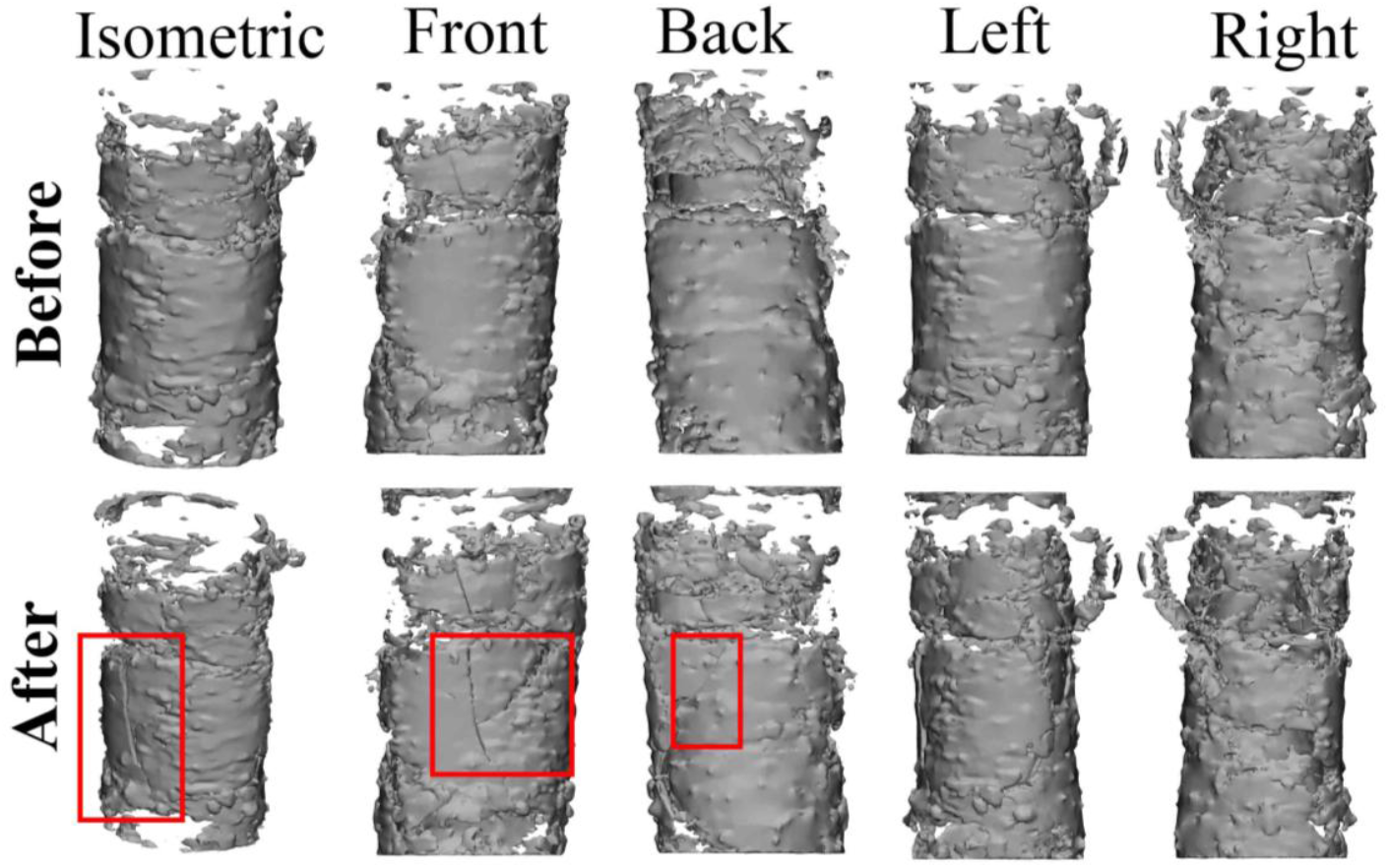
3D view of cylindrical calcification for sample #3 before and after angioplasty. Crack growth and nucleation is indicated with red boxed regions.

Figure 7 demonstrates representative histologic transverse VVG-stained images of the control (undamaged) segment and the segment from the angioplasty region (damaged) in lightly calcified specimens. Histologic evaluation of severely calcified tissues was inconclusive because specimens had to be decalcified prior to embedding in paraffin in order to avoid tearing when cut with a microtome. Angioplasty-induced damage manifested primarily as delamination along the internal elastic lamina (IEL), but tearing within the tunica media was also present. Both likely contributed to the formation of dissections seen on histology slides (Figure 7) and 3D reconstructions (Figure 2). The healthier arterial wall appeared overstretched after angioplasty, and remnant strains suggested appreciable damage inflicted on it by the balloon.

**Figure 7.**
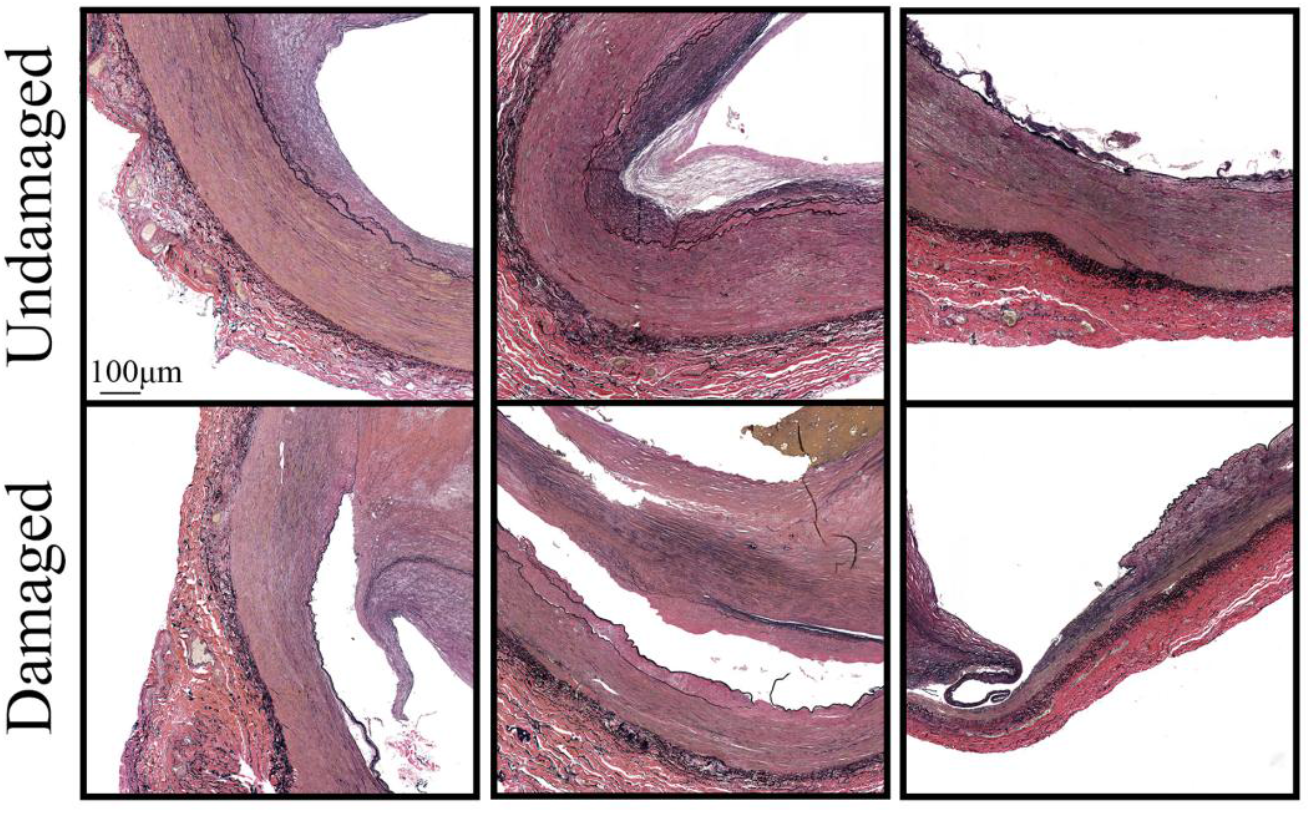
Verhoeff-Van Gieson (VVG)-stained transverse sections from sample #10 (left), #13 (middle), and #11 (right). All three samples had light calcification. The top images correspond to the control segment (undamaged), while the bottom images show segments from the angioplasty region. Note the delamination along the internal elastic lamina from both intimal and medial sides, tearing within the tunica media, and overstretching of the arterial wall as a result of angioplasty.

Experimental biaxial Cauchy stress-stretch relations for all tested specimens are summarized in Figure 8. Arteries are grouped by the calcium burden: red – severe and green – mild calcification. In severely calcified arteries, damage initiated on average at 1.11 ± 0.02 stretch and 44 ± 22 kPa stress longitudinally, and at 1.11 ± 0.03 stretch and 66 ± 27 kPa stress circumferentially. Mildly calcified specimens started accumulating damage at 1.15 ± 0.07 stretch and 57 ± 31 kPa stress longitudinally, and at 1.14 ± 0.06 stretch and 69 ± 46 kPa stress circumferentially. In general, damage initiated at lower biaxial stretches and lower longitudinal stresses in severely calcified specimens compared with lightly calcified arteries, but larger calcium burdens required more circumferential stress to initiate damage.

**Figure 8.**
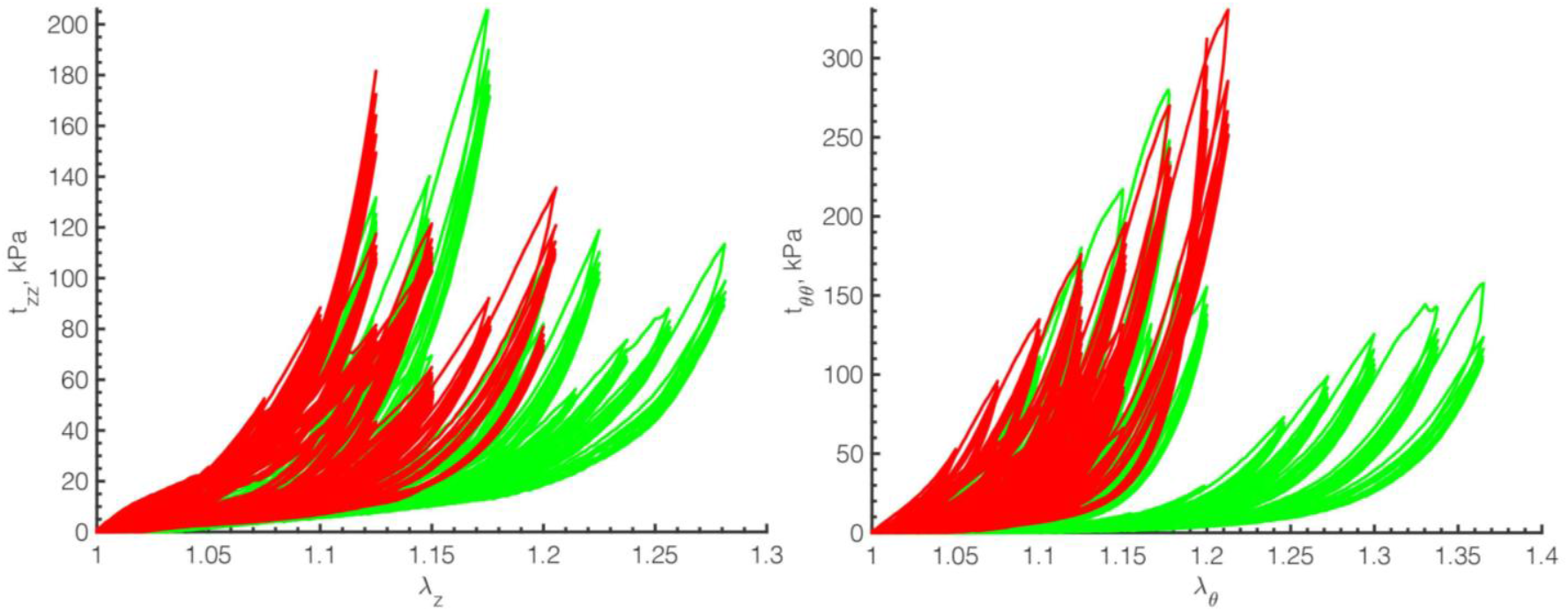
Cauchy stress-stretch relations in longitudinal (left, *z*) and circumferential (right, *θ*) directions demonstrating the biaxial assessment of damage in all 15 specimens. Colors represent arteries with severe (red) and mild (green) calcification. Note how arteries with larger calcium burdens are also stiffer, particularly in the circumferential direction.

Two reconstructed fluoroscopic views of sample #4 after angioplasty from two different viewing rotational angles are presented in Figure 9. Fluoroscopic view #1 shows a relatively open lumen with calcification spanning part of the arterial wall. Fluoroscopic view #2 shows a rotated view with a substantially obstructed lumen with heavy calcification spanning nearly the entire length of the sample. Recreated digital subtracted angiography (DSA) images for the same fluoroscopic views are also presented, also showing what appears to be an open lumen in view #1 and a still occluded lumen in view #2.

**Figure 9.**
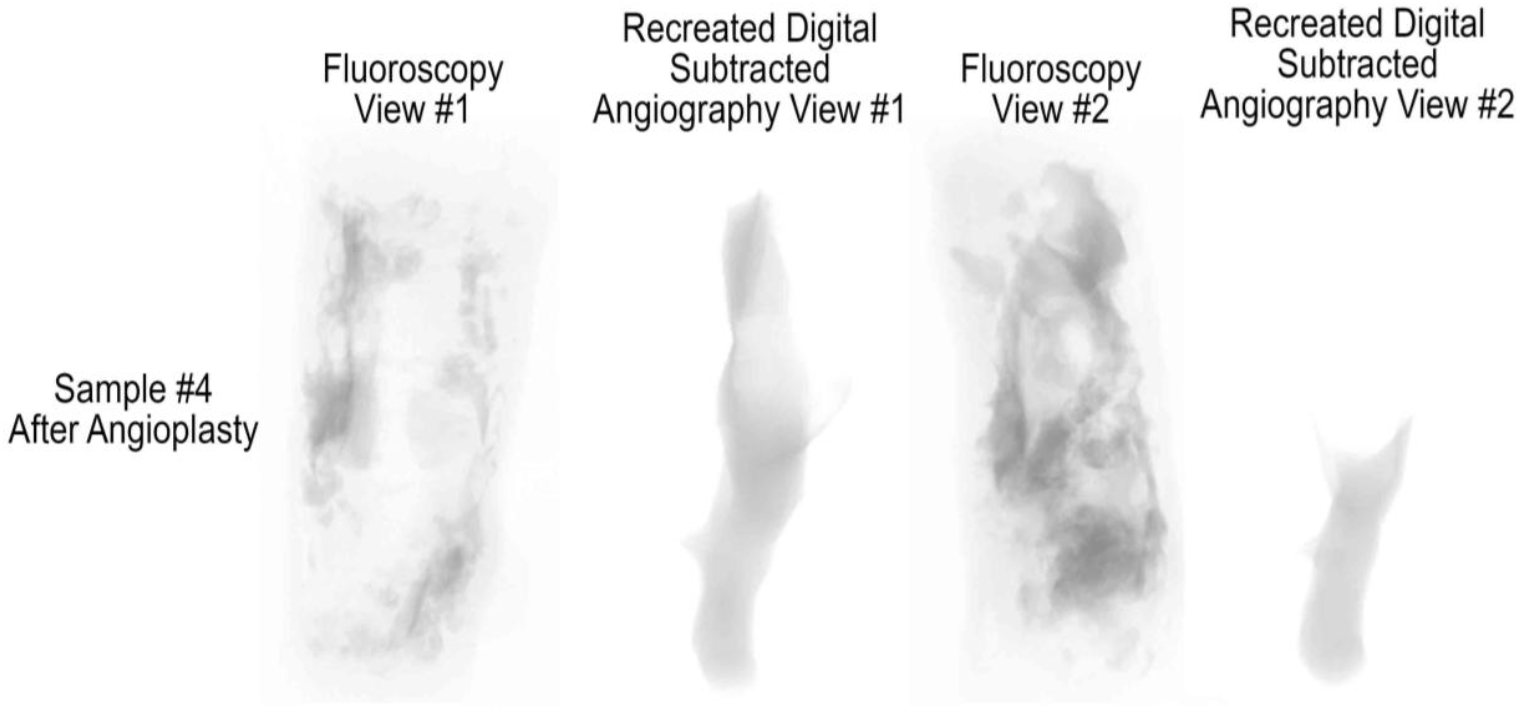
Reconstructed fluoroscopic views and recreated digital subtracted angiography of sample #4 using Mimics from two different rotational angles.

## 4. DISCUSSION

Balloon angioplasty is one of the most common options for treating symptomatic PAD, but its clinical outcomes continue to disappoint, particularly when managing highly calcified lesions^28^. Although the association between the high calcium burden and the inferior outcomes is well known^29^, the exact reasons are not completely understood. In this study we have used high-resolution CT imaging, structural analysis, and mechanical testing to determine what happens to diseased human FPAs during balloon angioplasty. Our data demonstrate that in severely calcified specimens, calcification typically manifests as calcium rings or large plate-like deposits that can span the entire circumference of the artery. In these specimens, damage caused by angioplasty resulted in the formation of longitudinal cracks. In samples with large, thick plate-like calcium deposits that did not span the entire circumference, little to no evidence of longitudinal cracking was present, but the luminal gains were higher. Clinical studies suggest that in addition to serving as a mechanical barrier to angioplasty, calcification also inhibits drug absorption^30^, which may contribute to the inferior results^31,32^ of drug-coated angioplasty balloons^33^ that have initially promised improved patency. A recent study^30^ found that calcifications spanning the entire circumference of the artery represented a major barrier to drug absorption, which in combination with more than modest luminal gains observed here could explain the high reintervention rate in these arteries. Adjuncts to balloon angioplasty, such as intravascular lithotripsy or atherectomy, may potentially reduce the effects of severe calcification, although their safety and efficiency for treating PAD lesions still need to be fully understood. In addition, it is likely that stenting following angioplasty would continue to be required due to the high prevalence of flow-limiting arterial dissections in both severely and non-severely calcified lesions.

Specifically, our data demonstrate that dissections were present in more than half of the samples after angioplasty, and the higher prevalence (75%) was observed in severely calcified arteries. These findings highlight the importance of evaluating the lesion post angioplasty. Dissections can compromise patency and increase the risk of distal embolization, and a recent study investigating the angiographic dissection patterns in superficial femoral artery lesions reported a prevalence of 42% in a retrospective, multicenter analysis of 621 patients^11^. Clinical evidence suggests that the use of longer balloons may help prevent dissections in chronic FPA obstructions^21^. Our results obtained with short 2 cm balloons illustrate the squeezing effect near the end of the balloon (i.e., #10, 12, and 15 in Figure 2), which in some cases have led to dissection. Non-uniform load distribution, in combination with the heterogeneous structure of the diseased FPA wall, likely contributes to the development and progression of dissections at or near the balloon ends. Time of inflation is another parameter that can contribute to the formation of the dissection. Our balloons were inflated for approximately 30 minutes to allow for the specimen to be scanned in the micro-CT, and a recent study that evaluated the effects of prolonged angioplasty on the formation of FPA dissections found that ballooning for >3 minutes was effective in preventing severe dissections^34^. While our *ex vivo* data cannot be directly compared with these *in vivo* results, these data suggest that shorter inflation times could potentially be associated with an even higher prevalence of arterial dissections than found by our current analysis. Importantly, though we have observed little to no arterial debris in the 3D printed fixtures after angioplasty, the presence of physiologic flow in conjunction with a significant number of observed dissections do not exclude the possibility of distal embolization upon balloon deflation and require further analysis.

Histological evaluation allowed us to better understand the characteristics of damage caused by the angioplasty balloon on the FPA wall. Our results demonstrate that damage manifested primarily as a delamination along the IEL interface and tearing within the tunica media layer, which agrees with prior findings obtained using supraphysiologic planar biaxial extension^27^. In the FPA, the IEL has a form of an elastic sheet (sometimes a double sheet) that acts as a barrier between the endothelial cells of the tunica intima and the smooth muscle cells (SMCs) of the tunica media^25,35^. SMCs are connected to the IEL via radially oriented elastin protrusions and oxytalan fibers, which facilitate mechanotransduction^36,37^. Degradation and fragmentation of these fibers due to age, disease, and proteolytic activity, may result in weaker interfacial links at the cell-elastic laminae boundary, thus resulting in the delamination along the IEL interface observed here and in other studies^37^. Additionally, older and more diseased FPAs tend to exhibit more medial degeneration and SMC loss compared to younger and healthier arteries in which circumferentially-oriented SMCs may act as a reinforcement^38–40^. Older arteries also accumulate more proteoglycans and glycosaminoglycans with age^26^ that can increase Donnan swelling pressure and also contribute to easier delamination^37^.

Structural changes to the arterial wall are heavily influenced by the systemic risk factors, with age, smoking, and diabetes playing significant roles in PAD progression^41,42^. While the detailed analysis in the context of risk factors requires a much larger sample size, our current data were obtained for older (average age 69 ± 9 years), overweight (average BMI 31.2 ± 7.2) subjects, nearly half of whom had DM, and all but one were either current or former smokers. Many studies reported the effects of these risk factors on cardiovascular health. For example, both current and former smokers were found to be at a greater risk for suffering severe cardiovascular events, including the development of atherosclerosis and DM^43–46^. DM was associated^40^ with the presence of severe, ring-shaped, and plate-like arterial calcification patterns, which contributed to circumferential mechanical stiffening to the point that middle-aged diabetic FPAs acquired the circumferential stiffness of much older arteries^47^. In line with our current findings are previous reports suggesting that age, body mass index (BMI), and DM had significant effects on damage experienced by human FPAs when stretched beyond their physiologic range^27^. In that study, damage initiation stretches and stresses were reported to reduce with age, subjects with higher BMI were shown to accumulate damage at lower stretches both longitudinally and circumferentially, and subjects with DM required higher stresses to begin accumulating damage circumferentially. Our current results demonstrate a similar trend whereby arteries from old and overweight subjects required lower biaxial stretches (1.13 ± 0.05) and stresses (50 ± 27 kPa longitudinally and 67 ± 36 kPa circumferentially) to begin damage accumulation. Also, in line with previous reports^27^, we have found that specimens with DM required higher circumferential stresses (92 ± 38 kPa) to induce damage as compared with arteries that did not have DM (45 ± 14 kPa), and our current results explain this by the presence of circumferential ring-shaped structures, aligning with the orientation of SMCs, that reinforce the artery around its circumference and contribute to its greater stiffness^47^.

While the exact mechanism for ring-like calcium formation remains to be further understood, it was previously hypothesized that under the influence of elevated glucose, a subset of SMCs within the tunica media lose their ability to express SMC-specific markers and start expressing osteogenic or chondrogenic markers^48,49^, thus transforming parts of the arterial wall into rings of bone tissue. As demonstrated in our current study, balloon angioplasty may fracture these rings and cause the cracks to propagate longitudinally, but it remains to be understood whether these cracks fuse together over time under the continued influence of hyperglycemia. Interestingly, we have also found that balloon angioplasty appeared to somewhat shift the calcification in the longitudinal direction, thereby stretching the tissue axially. While, as far as we know, this has not yet been reported in the FPA, similar results were described for the circumferentially calcified carotid artery plaques after balloon angioplasty^50^. Furthermore, axial tissue overstretch by the balloon and then by the deployed stent, was associated with the higher prevalence of restenosis after angioplasty in these arteries^50^. Our current results demonstrate that severely calcified FPAs start accumulating damage just after 1.11 biaxial stretch, but most stents that are used to treat PAD are usually oversized by at least 1.15, creating a chronic trauma that continues to stretch the diseased vessel beyond its elastic limit long after the patient leaves the operating room.

Reconstructed fluoroscopic and digital subtraction angiography views obtained from two rotational angles suggested a relatively patent lumen, consistent with an apparently successful balloon angioplasty. However, imaging from an alternative angle revealed persistent, high-grade stenosis with substantial residual calcification. This discrepancy underscores the critical importance of incorporating intravascular ultrasound (IVUS) when assessing lesion morphology and angioplasty outcomes. Because fluoroscopic appearances can vary markedly with imaging orientation, IVUS provides essential cross-sectional detail that can prevent misinterpretation and ultimately improve patency outcomes following endovascular intervention.

While the presented results allow a better understanding of the effects of balloon angioplasty on human FPAs with different calcium burdens, they should be viewed in the context of study limitations. First and foremost, we have reported *in vitro* results that may not directly translate to *in vivo* scenarios that include physiologic flow, the influence of surrounding tissues and tethering branches, the response of live cells, and circulating factors. Unfortunately, limitations of the clinical imaging technology currently do not allow to obtain sufficiently high-resolution images to perform this study *in vivo*, but promising new techniques utilizing machine learning are being developed to circumvent some of these issues. The second limitation is the setup we have used to obtain our high-resolution CTs. Due to similar attenuation coefficients between the arterial wall and saline, it is challenging to scan the artery submerged. While 30min required for our scans have not resulted in drying out of the tissue, scanning longer segments may need additional treatments to prevent this effect from happening. Lastly, our study was limited to 15 specimens with different calcification burdens, but a larger sample size is needed to better understand the effects of risk factors and calcification patterns on angioplasty results and perform proper statistical comparisons. We hope to address these limitations in our future studies to better understand the causes of poor clinical outcomes of endovascular PAD treatments.

## ACKNOWLEDGMENTS

This work was supported in part by National Institutes of Health under awards HL125736, HL180371, and P20GM152301. The authors wish to acknowledge Live On Nebraska for their help and support and thank tissue donors and their families for making this study possible.

